## Supplemental Figures for "Strong Ethanol- and Frequency-Dependent Ecological Interactions in a Community of Wine-Fermenting Yeasts"

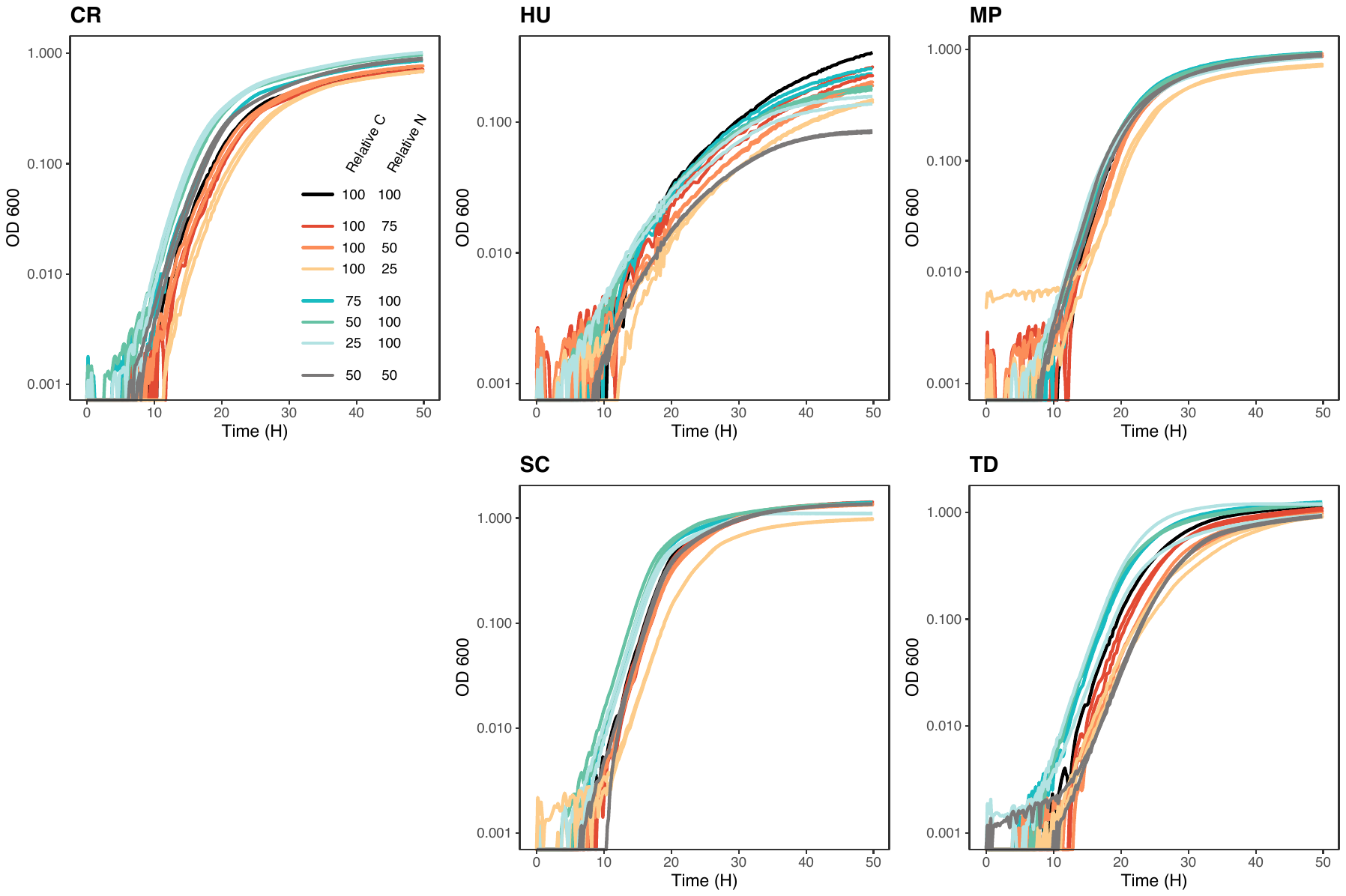

**Figure S1: Growth rates in a synthetic grape juice like medium are barely affected by removing up to 75% of the carbon or nitrogen sources in the medium.**

**Figure S2 (next page): Consensus interaction networks average the outcomes of three experimental replicates.** The blue stars in some Rep II networks indicate that the two initial conditions of the same pair did not reach the same outcome, with one condition reaching exclusion and one maintaining coexistence (at the fraction indicated by the blue star) at the end of the 7-day cycle. The gray arrows in the Rep II 5ABV network indicate that the coculture collapsed.

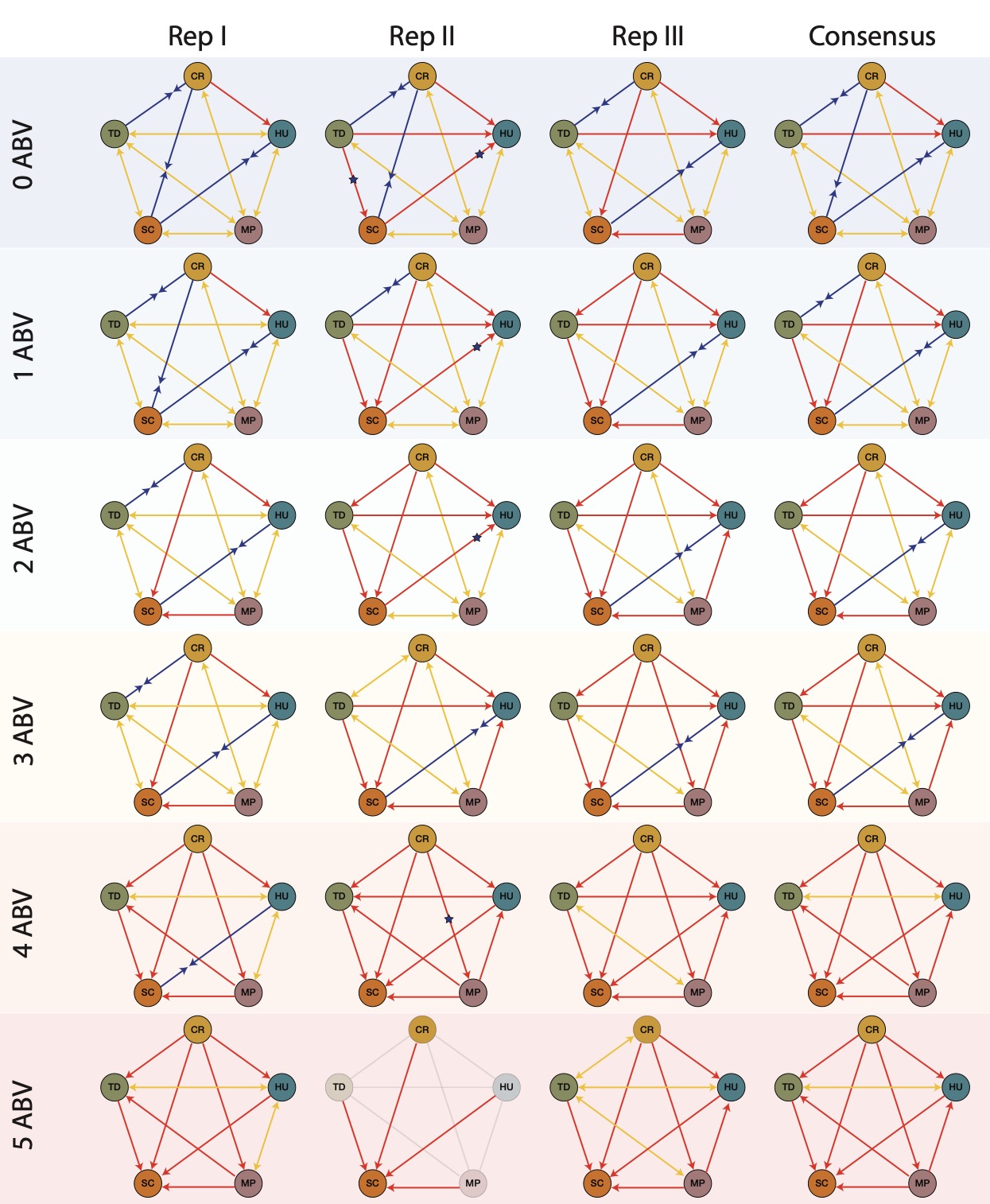
